# Admixture-enabled selection for rapid adaptive evolution in the Americas

**DOI:** 10.1101/783845

**Authors:** Emily T. Norris, Lavanya Rishishwar, Aroon T. Chande, Andrew B. Conley, Kaixiong Ye, Augusto Valderrama-Aguirre, I. King Jordan

**Affiliations:** School of Biological Sciences, Georgia Institute of Technology, Atlanta, GA, USA; IHRC-Georgia Tech Applied Bioinformatics Laboratory, Atlanta, GA, USA; PanAmerican Bioinformatics Institute, Cali, Valle del Cauca, Colombia; Department of Genetics, University of Georgia, Athens, GA, USA; Institute of Bioinformatics, University of Georgia, Athens, GA, USA; Biomedical Research Institute (COL0082529), Cali, Colombia; Universidad Santiago de Cali, Cali, Colombia

**Keywords:** Rapid adaptive evolution, positive selection, genetic ancestry, admixture, population genomics, polygenic traits

## Abstract

**Background:** Admixture occurs when previously isolated populations come together and exchange genetic material. We hypothesized that admixture can enable rapid adaptive evolution in human populations by introducing novel genetic variants (haplotypes) at intermediate frequencies, and we tested this hypothesis via the analysis of whole genome sequences sampled from admixed Latin American populations in Colombia, Mexico, Peru, and Puerto Rico.

**Results:** Our screen for admixture-enabled selection relies on the identification of loci that contain more or less ancestry from a given source population than would be expected given the genome-wide ancestry frequencies. We employed a combined evidence approach to evaluate levels of ancestry enrichment at (1) single loci across multiple populations and (2) multiple loci that function together to encode polygenic traits. We found cross-population signals of African ancestry enrichment at the major histocompatibility locus on chromosome 6, consistent with admixture-enabled selection for enhanced adaptive immune response. Several of the human leukocyte antigen genes at this locus (*HLA-A*, *HLA-DRB51* and *HLA-DRB5*) showed independent evidence of positive selection prior to admixture, based on extended haplotype homozygosity in African populations. A number of traits related to inflammation, blood metabolites, and both the innate and adaptive immune system showed evidence of admixture-enabled polygenic selection in Latin American populations.

**Conclusions:** The results reported here, considered together with the ubiquity of admixture in human evolution, suggest that admixture serves as a fundamental mechanism that drives rapid adaptive evolution in human populations.

## Background

Admixture is increasingly recognized as a ubiquitous feature of human evolution [1]. Recent studies on ancient DNA have underscored the extent to which human evolution has been characterized by recurrent episodes of population isolation and divergence followed by convergence and admixture. In this study, we considered the implications of admixture for human adaptive evolution [2]. We hypothesized that admixture is a critical mechanism that enables rapid adaptive evolution in human populations, and we tested this hypothesis via the analysis of admixed genome sequences from four Latin American populations: Colombia, Mexico, Peru, and Puerto Rico. We refer to the process whereby the presence of distinct ancestry-specific haplotypes on a shared population genomic background facilitates adaptive evolution as ‘admixture-enabled selection’.

The conquest and colonization of the Americas represents a major upheaval in the global migration of our species and is one of the most abrupt and massive admixture events known to have occurred in human evolution [3, 4]. The ancestral source populations – from Africa, Europe, and the Americas – that admixed to form modern Latin American populations evolved separately for tens-of-thousands of years before coming together over the last 500 years. This 500-year time frame, corresponding to approximately 20 generations, amounts to less than 1% of the time that has elapsed since modern humans first emerged from Africa [5, 6]. Considered together, these facts point to admixed Latin American populations as an ideal system to study the effects of admixture on rapid adaptive evolution in humans [7].

A number of previous studies have considered the possibility of admixture-enabled selection in the Americas, yielding conflicting results. On the one hand, independent studies have turned up evidence for admixture-enabled selection at the major histocompatibility complex (MHC) locus in Puerto Rico [8], Colombia [9], and Mexico [10], and another study found evidence for admixture-enabled selection on immune system signaling in African-Americans, particularly as it relates to influenza and malaria response [11]. Together, these studies highlighted the importance of the immune system as a target for admixture-enabled selection among a diverse group of admixed American populations. However, a follow up study on a different cohort of African-Americans found no evidence for admixture-enabled selection in the Americas [12]. The latter study concluded that the observed differences in local ancestry reported by previous studies, which were taken as evidence for selection, could have occurred by chance alone given the large number of hypotheses that were tested (i.e. the number of loci analyzed across the genome). This work underscored the importance of controlling for multiple hypothesis testing when investigating the possibility of admixture-enabled selection in the Americas.

We attempted to resolve this conundrum by performing integrated analyses that combine information from (1) single loci across multiple populations and (2) multiple loci that encode polygenic traits. We also used admixture simulation, along with additional lines of evidence from haplotype-based selection scans, to increase the stringency of, and confidence in, our screen for admixture-enabled selection. This combined evidence approach has proven to be effective for the discovery of admixture-enabled selection among diverse African populations [13, 14]. We found evidence for admixture-enabled selection at the MHC locus across multiple Latin American populations, consistent with previous results, and our polygenic screen uncovered novel evidence for adaptive evolution on a number of inflammation, blood, and immune related traits.

## Results

### Genetic ancestry and admixture in Latin America

We inferred patterns of genetic ancestry and admixture for four Latin American (LA) populations characterized as part of the 1000 Genomes Project: Colombia (*n*=94), Mexico (*n*=64), Peru (*n*=85), and Puerto Rico (*n*=104) (Figure 1). Genome-wide continental ancestry fractions were inferred using the program ADMIXTURE [15], and local (haplotype-specific) ancestry was inferred using the program RFMix [16]. The results from both programs are highly concordant (Additional file 1: Figure S1). As expected [17–20], the four LA populations show genetic ancestry contributions from African, European, and Native American source populations, and they are distinguished by the relative proportions of each ancestry. Overall, these populations show primarily European ancestry followed by Native American and African components. Puerto Rico has the highest European ancestry, whereas Peru shows the highest Native American ancestry. Mexico shows relatively even levels of Native American and European ancestry, while Colombia shows the highest levels of three-way admixture. Individual genomes vary greatly with respect to the genome-wide patterns of local ancestry, i.e. the chromosomal locations of ancestry-specific haplotypes (Additional file 1: Figure S2). If the process of admixture is largely neutral, then we expect ancestry-specific haplotypes to be randomly distributed throughout the genome in proportions corresponding to the genome-wide ancestry fractions.

**Figure 1.**
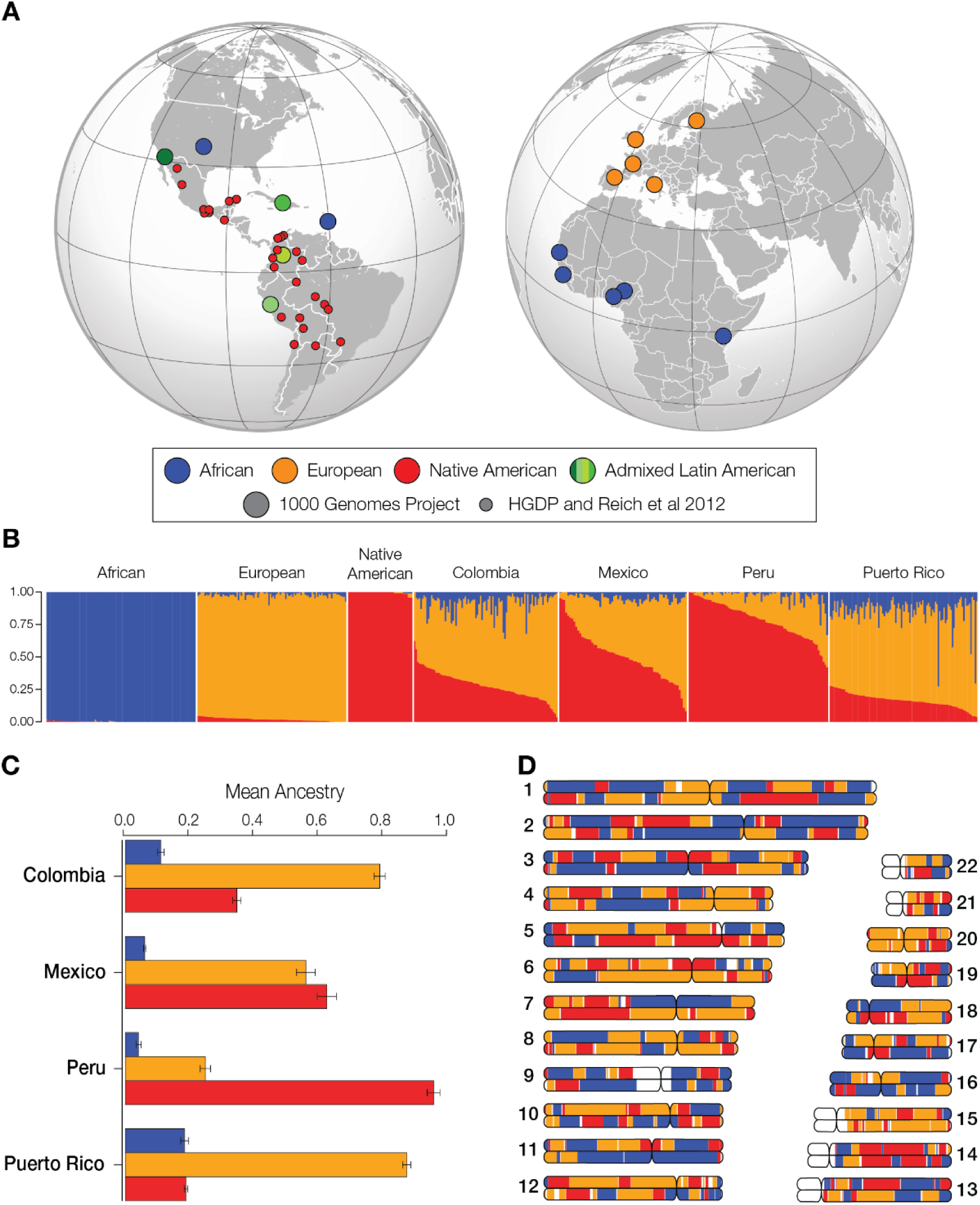
Genetic ancestry and admixture in Latin America. (A) The global locations of the four LA populations analyzed here (green) are shown along with the locations of the African (blue), European (orange), and Native American (red) reference populations. The sources of the genomic data are indicated in the key. (B) ADMIXTURE plot showing the three-way continental ancestry components for individuals from the four LA populations – Colombia, Mexico, Peru, and Puerto Rico – compared to global reference populations. (C) The mean (±*se*) continental ancestry fractions for the four LA populations. (D) Chromosome painting showing the genomic locations of ancestry-specific haplotypes for an admixed LA genome.

### Ancestry enrichment and admixture-enabled selection

For each of the four LA populations, local ancestry patterns were used to search for specific loci that show contributions from one of the three ancestral source populations which are greater than can be expected based on the genome-wide ancestry proportions for the entire population (Additional file 1: Figure S3). The ancestry enrichment metric that we use for this screen (*z_anc_*) is expressed as the number of standard deviations above or below the genome-wide ancestry fraction. Previous studies have used this general approach to look for evidence of admixture-enabled selection at individual genes within specific populations, yielding mixed results [8–12]. For this study, we have added two new dimensions to this general approach in an effort to simultaneously increase the confidence for admixture-enabled selection inferences and to broaden the functional scope of previous studies. To achieve these ends, we searched for (1) concordant signals of ancestry enrichment for single genes (loci) across multiple populations, and (2) concordant signals of ancestry enrichment across multiple genes that function together to encode polygenic phenotypes. The first approach can be considered to increase specificity, whereas the second approach increases sensitivity. Loci that showed evidence for ancestry enrichment using this combined approach were interrogated for signals of positive selection using the integrated haplotype score (iHS) [21] to further narrow the list of potential targets of admixture-enabled selection.

### Single gene admixture-enabled selection

Gene-specific ancestry enrichment values (*z_anc_*) were computed for each of the three continental ancestry components within each of the four admixed LA populations analyzed here. We then integrated gene-specific *z_anc_* values across the four LA populations using a Fisher combined score (*F*_*CS*_). The strongest signals of single gene ancestry enrichment were seen for African ancestry at the major histocompatibility complex (MHC) locus on the short arm of chromosome 6 (Figure 2A). Three out of the four LA populations show relatively high and constant ancestry enrichment across this locus, with the highest levels of enrichment seen for Mexico and Colombia (Figure 2B). This signal is robust to control for multiple statistical tests using the Benjamini–Hochberg false discovery rate (FDR).

**Figure 2.**
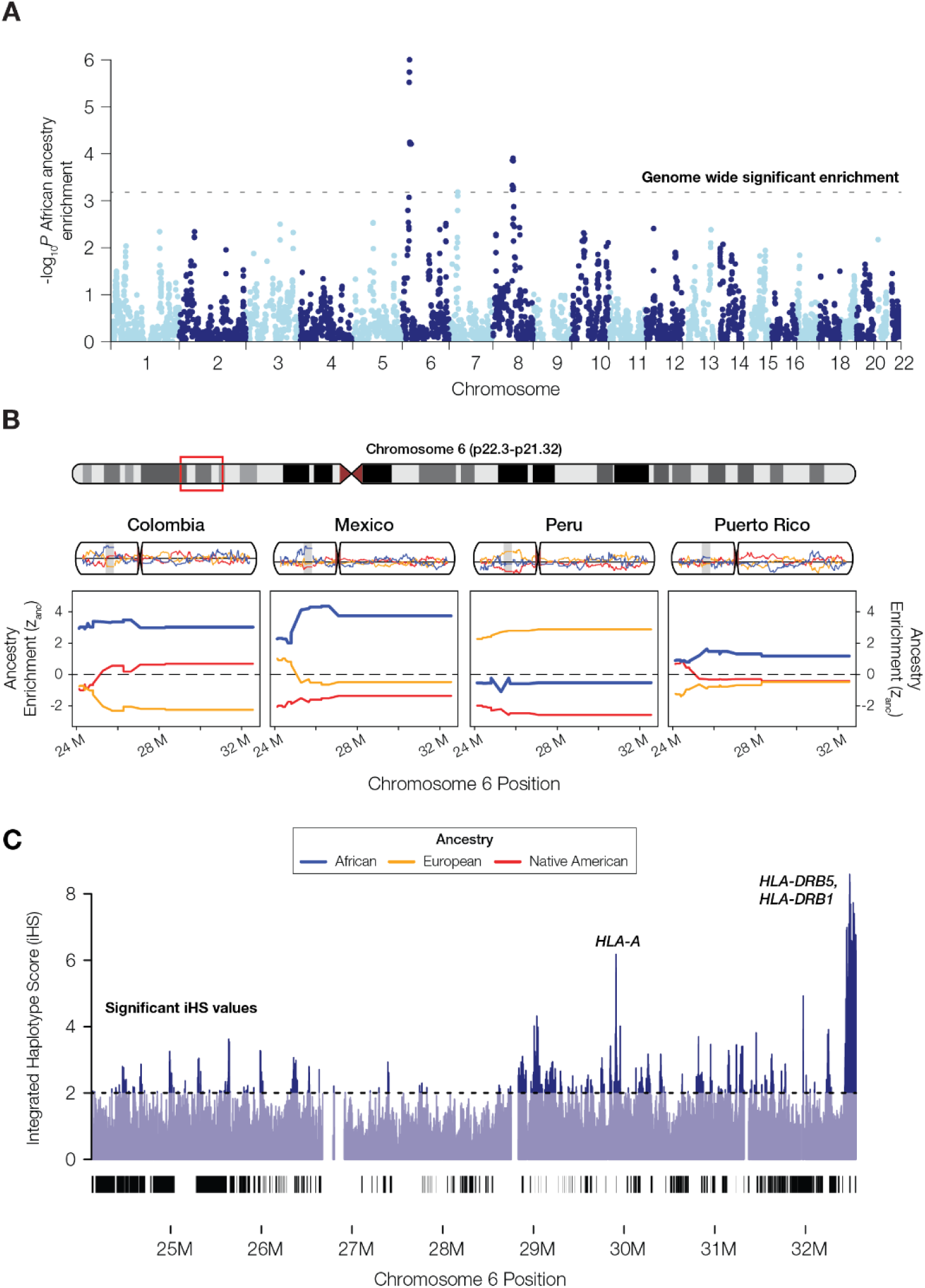
African ancestry enrichment at the major histocompatibility complex (MHC) locus. (A) Manhattan plot showing the statistical significance of African ancestry enrichment across the genome. (B) Haplotype on chromosome 6 with significant African ancestry enrichment for three of the four LA populations: Colombia, Mexico, and Puerto Rico. This region corresponds to the largest peak of African ancestry enrichment on chromosome 6 seen in panel A. Population-specific African (blue), European (orange), and Native American (red) ancestry enrichment values (*z_anc_*) are shown for chromosome 6 and the MHC locus. (C) Integrated haplotype score (iHS) values for African continental population from the 1KGP are shown for the MHC locus; peaks correspond to putative positively selected human leukocyte antigen (*HLA*) genes.

We simulated random admixture across the four LA populations, parameterized by their genome-wide ancestry proportions, to further assess the probability that this signal could be generated by chance alone (i.e. by genetic drift). Based on this simulation, the observed levels of cross-population African ancestry enrichment at the MHC locus are highly unlikely to have occurred by chance (*P* < 5 × 10^−5^), whereas the observed patterns of European and Native American ancestry enrichment are consistent with the range of expected levels generated by the random admixture simulation (Additional file 1: Figure S4). Results of the admixture simulation analysis were also used to demonstrate that the cross-population approach to single locus ancestry-enrichment is sufficiently powered to detect selection at the population sizes analyzed here (Additional file 1: Figures S5 and S6). The statistical power of the ancestry enrichment approach used in this study rests on the cross-population comparisons, as the probability of observing the same ancestry enrichment at the same locus across multiple LA populations is diminishingly low.

The MHC locus of chromosome 6 also shows a number of peaks for the iHS metric of positive selection from the African continental population (Figure 2C). These peaks rise well above the value of 2, which is taken as a threshold for putative evidence of positive selection [21]. The highest African iHS scores are seen for the human leukocyte antigen (HLA) encoding genes *HLA-A*, *HLA-DRB5*, and *HLA-DRB1* (Figure 3A and 3B). These HLA protein encoding genes make up part of the MHC class I (*HLA-A*) and MHC class II (*HLA-DRB5*, and *HLA-DRB1*) antigen presenting pathways of the adaptive immune system (Figure 3C), consistent with shared selective pressures on immune response in admixed LA populations.

**Figure 3.**
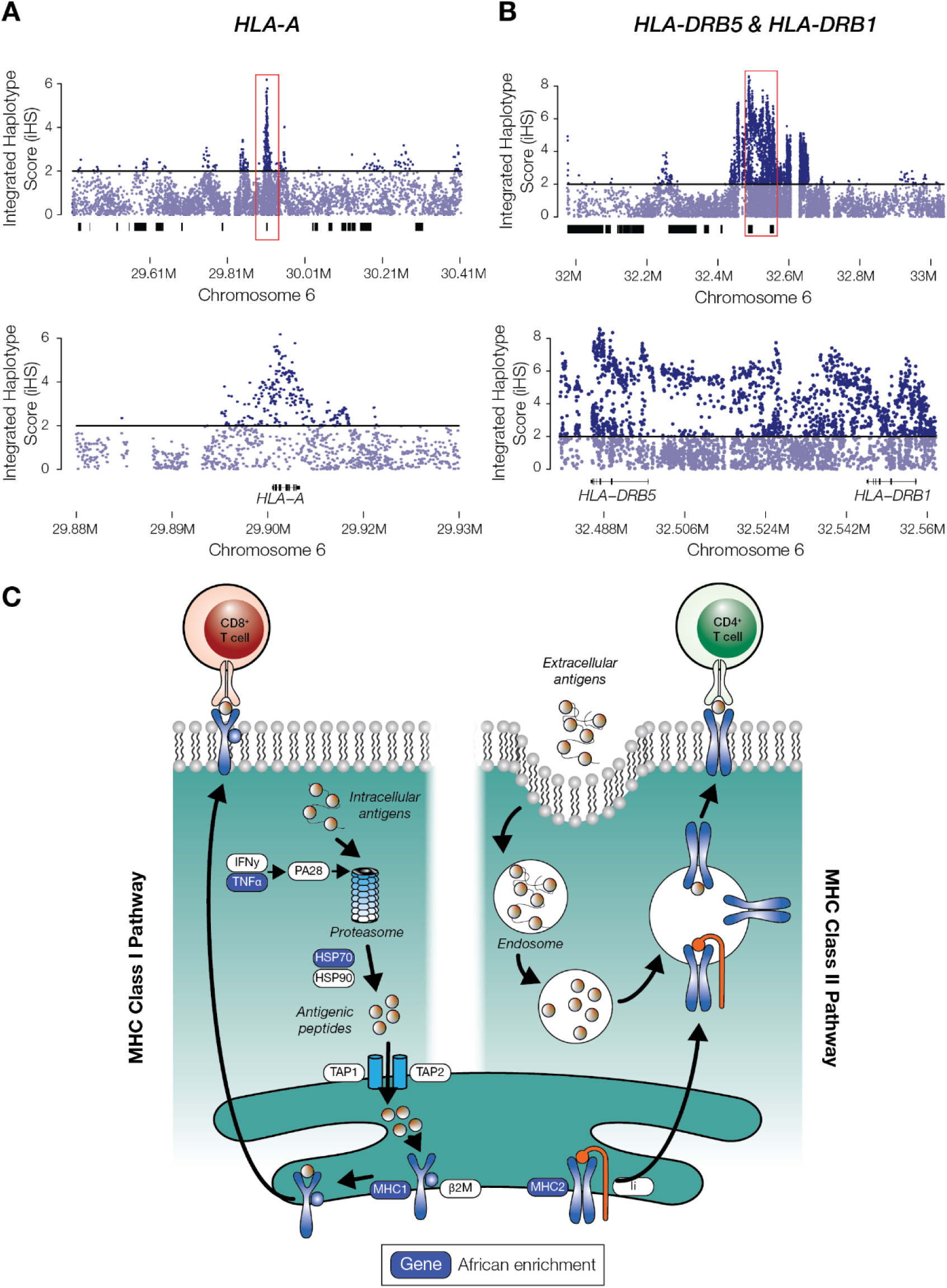
Admixture-enabled selection at human leukocyte antigen (*HLA*) genes. Integrated haplotype score (iHS) peaks for the African continental population from the 1KGP are shown for (A) the MHC Class I gene *HLA-A* and (B) the MHC Class II genes *HLA-DRB5* and *HLA-DRB1*. (C) Illustration of the MHC Class I and MHC Class II antigen presenting pathways, with African enriched genes shown in blue.

We modeled the magnitude of selection pressure that would be needed to generate the observed levels of cross-population African ancestry enrichment at the MHC locus, using a tri-allelic recursive population genetics model that treats ancestry haplotype fractions as allele frequencies (Figure 4). The average selection coefficient value for African MHC haplotypes is *s*=0.05 (Additional file 1: Figure S7), indicating strong selection at this locus over the last several hundred years since the admixed LA populations were formed, consistent with previous work [10].

**Figure 4.**
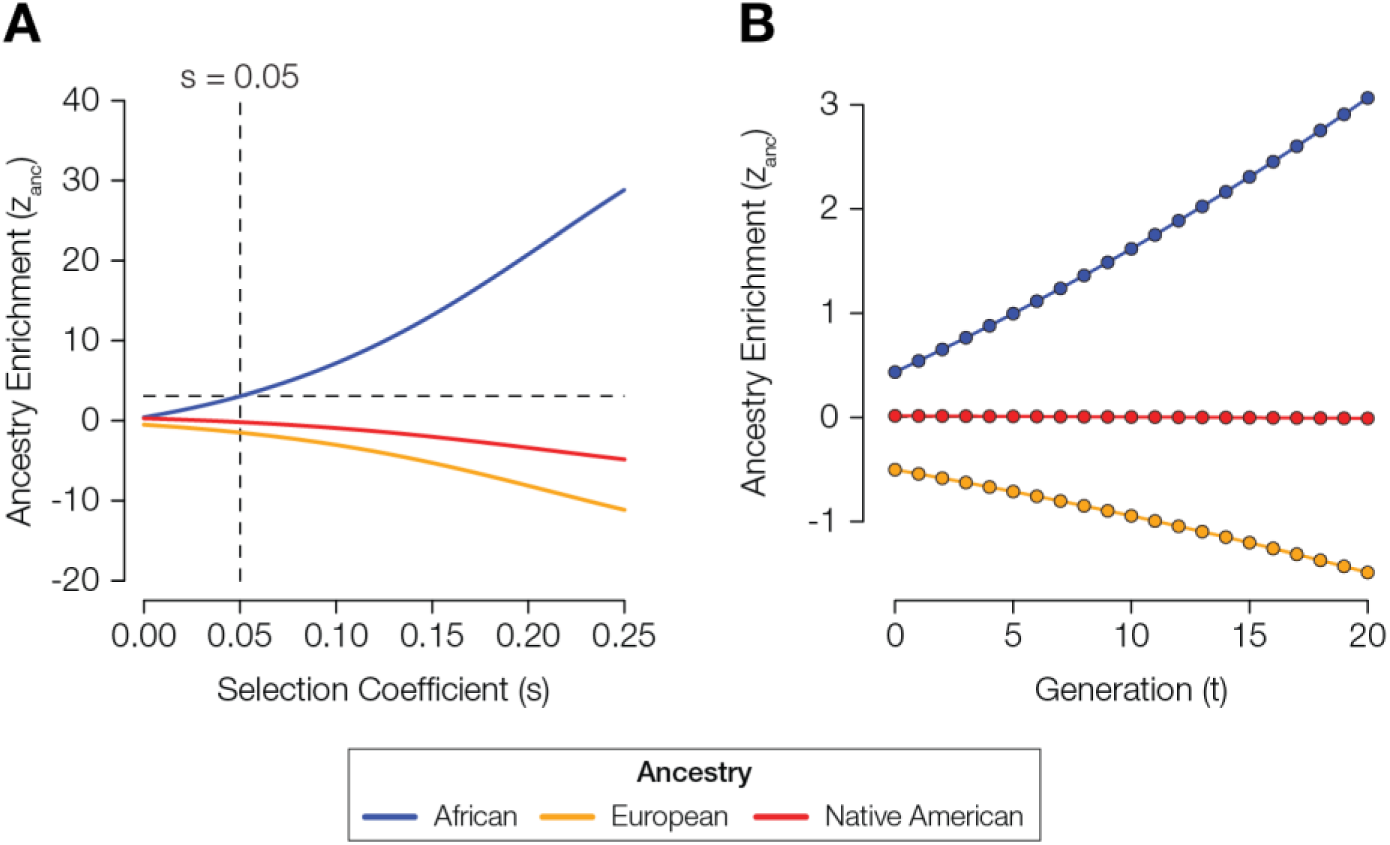
Model of ancestry-enabled selection at the MHC locus in the Colombia population. (A) Modeled levels of ancestry enrichment and depletion (*z_anc_*, y-axis) corresponding to a range of different selection coefficients (*s*, x-axis): African (blue), European (orange), and Native American (red). The intersection of the observed level of African ancestry enrichment at the MHC locus and the corresponding *s*-value is indicated with dashed lines. (B) The trajectory of predicted ancestry enrichment and depletion (*z_anc_*, y-axis) over time (*t* generations, x-axis) is shown for the inferred selection coefficient of *s*=0.05.

### Polygenic admixture-enabled selection

For each of the three continental ancestry components, we combined gene-specific ancestry enrichment values (*z_anc_*), for genes that function together to encode polygenic phenotypes, via the polygenic ancestry enrichment score (*PAE*) (Figure 5A). Observed *PAE* values were compared to expected values generated by randomly permuting size-matched gene sets to search for functions (traits) that show evidence of admixture-enabled selection (Additional file 1: Figure S8). As with the single locus approach, we narrowed our list of targets to traits that showed evidence of polygenic admixture enrichment across multiple LA populations. This approach yielded evidence of statistically significant enrichment and depletion, across multiple ancestries, for a number of inflammation, blood, and immune related traits (Figure 5B). Inflammation related phenotypes that show polygenic ancestry enrichment include a variety of skin conditions and rheumatoid arthritis. A number of different blood metabolite pathways show evidence for primarily European and Native American ancestry enrichment, while both the adaptive and innate components of the immune system show evidence of admixture-enabled selection.

**Figure 5.**
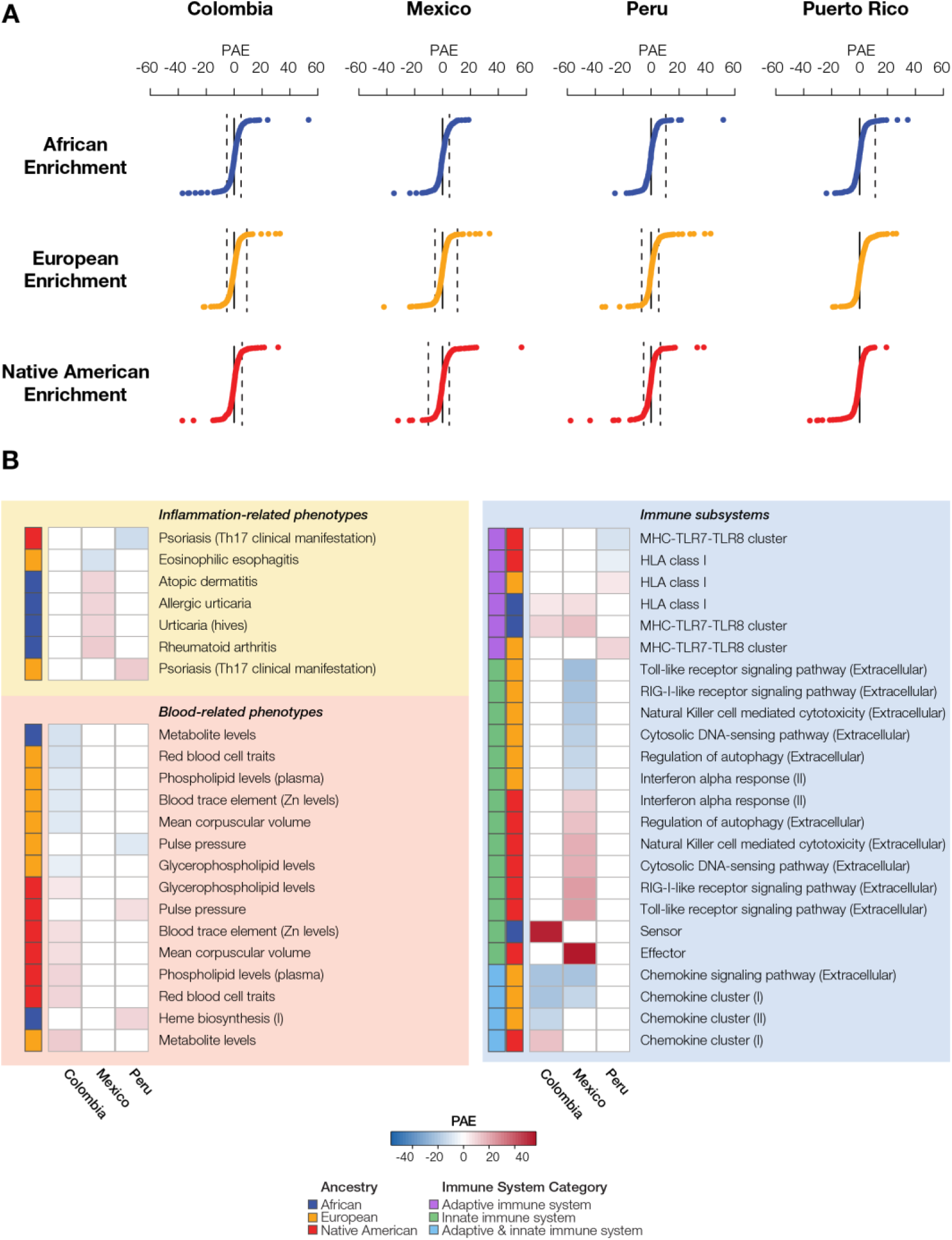
Polygenic ancestry enrichment (*PAE*) and admixture-enabled selection. (A) Distributions of the *PAE* test statistic are shown for each of the three ancestry components – African (blue), European (orange), and Native American (red) – across the four LA populations. Points beyond the dashed lines correspond to polygenic traits with statistically significant *PAE* values, after correction for multiple tests. (B) Polygenic traits that show evidence of *PAE* in multiple LA populations. *PAE* values are color coded as shown in the key, and the ancestry components are indicated for each trait. Immune system traits are divided into adaptive (purple), innate (green), or both (blue).

Several interconnected pathways of the innate immune system – the RIG-I-like receptor signaling pathway, the Toll-like receptor signaling pathway, and the cytosolic DNA-sensing pathway – all show evidence of Native American ancestry enrichment (Figure 6). All three of these pathways are involved in rapid, first line immune response to a variety of RNA and DNA viruses as well as bacterial pathogens. Genes from these pathways that show evidence of Native American ancestry enrichment encode a number of distinct interferon, interleukin, and cytokine proteins.

**Figure 6.**
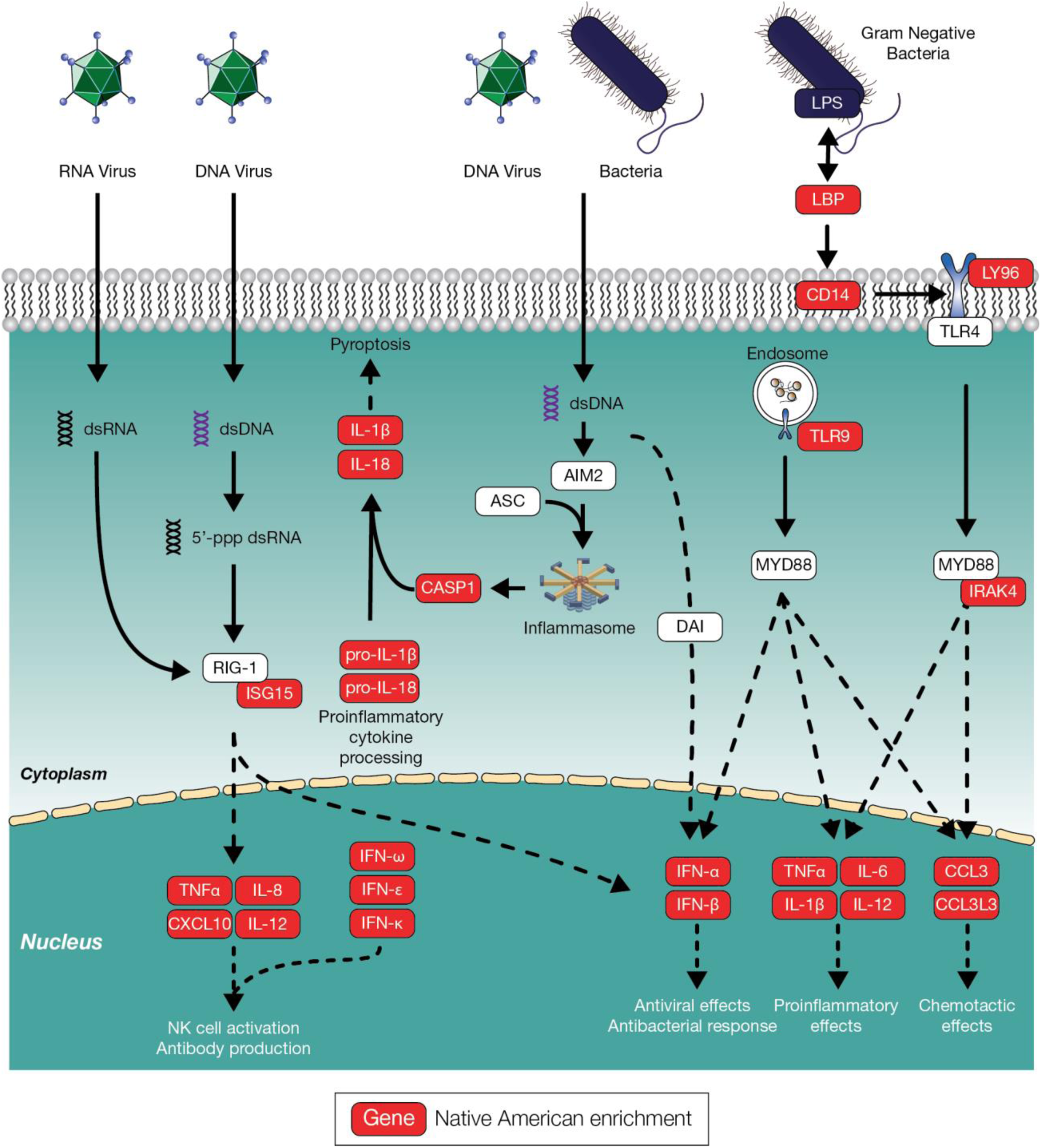
Innate immune system pathways showing Native American ancestry enrichment. Illustration of three interconnected pathways from the innate immune system – the RIG-I-like receptor signaling pathway, the Toll-like receptor signaling pathway, and the cytosolic DNA-sensing pathway – highlighting genes (proteins) that show Native American ancestry enrichment.

## Discussion

### Rapid adaptive evolution in humans

Human adaptive evolution is often considered to be a slow process, which is limited by relatively low effective population sizes and long generation times [22–24]. The rate of human adaptive evolution is further constrained by the introduction of new mutations [25]. Initially, positive selection acts very slowly to gradually increase the frequency of newly introduced beneficial mutations, which by definition are found at low population frequencies. The process of admixture, whereby previously diverged populations converge, brings together haplotypes that have not previously existed on the same population genomic background [26]. In so doing, it can provide raw material for rapid adaptive evolution in the form of novel variants that are introduced at intermediate frequencies, many of which may have pre-evolved adaptive utility [7].

### Admixture and rapid adaptive evolution

Our results suggest that admixture can enable extremely rapid adaptive evolution in human populations. In the case of the LA populations studied here, we found evidence of adaptive evolution within the last 500 years (or ∼20 generations) since the conquest and colonization of the Americas began [3, 4]. We propose that, given the ubiquity of admixture among previously diverged populations [1, 2], it should be considered as a fundamental mechanism for the acceleration of human evolution.

The haplotypes that show evidence of ancestry enrichment in our study evolved separately for tens-of-thousands of years in the ancestral source populations – African, European, and Native American – that mixed to form modern, cosmopolitan LA populations. Many of these haplotypes are likely to contain variants, or combinations of variants, that provided a selective advantage in their ancestral environments [27]. These adaptive variants would have increased in frequency over long periods of time and then later provided source material for rapid adaptation of admixed populations, depending on their utility in the New World environment. Variants that reached high frequency in ancestral source populations via genetic drift could also serve as targets for positive selection in light of the distinct environments and selection pressures faced by modern admixed populations. In either case, admixture-enabled selection can be taken as a special case of selection on standing variation, or soft selective sweeps, underscoring its ability to support rapid adaptation in the face of novel selective pressures [28, 29].

### Single locus versus polygenic selection

Our initial analysis of individual LA populations turned up numerous instances of apparent ancestry enrichment genome-wide, including enrichment for all three ancestry components in each of the four populations studied here (Additional file 2: Table S1). However, when ancestry enrichment signals were combined across all four populations, only a handful of significant results remained after correcting for multiple tests. Finally, when random admixture was simulated, only two peaks of African ancestry enrichment were found to be shared among populations at levels greater than expected by chance (Figure 2 and Additional file 1: Figure S4). These findings support the conservative nature of our combined evidence approach to using cross-population ancestry enrichment as a criterion for inferring admixture-enabled selection, and also reflect the fact that selection needs to be extremely strong to be detected at single loci. This is especially true given the relatively short period of time that has elapsed since modern LA populations were formed via admixture of ancestral source populations. The results of our population genetic model support this notion, showing an average selection coefficient value of *s*=0.05 for African haplotypes at the MHC locus.

A number of recent studies have underscored the ubiquity of polygenic selection on complex traits that are encoded by multiple genes, emphasizing the fact that weaker selection dispersed across multiple loci may be a more common mode of adaptive evolution than strong single locus selection [30–33]. The results of our polygenic ancestry enrichment analysis are consistent with these findings, as the polygenic approach yielded signals of admixture-enabled selection for numerous traits across different ancestry components and populations. Thus, the polygenic ancestry enrichment that we employed to infer admixture-enabled selection is both more biologically realistic and better powered compared to the single locus approach.

### Conclusions

We report abundant evidence for admixture-enabled selection within and between Latin American populations that were formed by admixture among diverse African, European, and Native American source populations within the last 500 years. The MHC locus shows evidence of particularly strong admixture-enabled selection for several *HLA* genes, all of which appear to contain pre-adapted variants that were selected prior to admixture in the Americas. In addition, a number of related immune system, inflammation and blood metabolite traits were found to evolve via polygenic admixture-enabled selection.

Over the last several years, it has become increasingly apparent that admixture is a ubiquitous feature of human evolution. Considering the results of our study together with the prevalence of admixture leads us to conclude that admixture-enabled selection has been a fundamental mechanism for driving rapid adaptive evolution in human populations.

## Methods

### Genomic data

Whole genome sequence data for four admixed LA populations – Colombia, Mexico, Peru, and Puerto Rico – were taken from the Phase 3 data release of the 1000 Genomes Project (1KGP) [34, 35]. Whole genome sequence data and whole genome genotypes for proxy ancestral reference populations from Africa, Asia, Europe, and the Americas were taken from multiple sources, including the 1KGP, the Human Genome Diversity Project (HGDP) [36] and a previous study on Native American genetic ancestry [37] (Additional file 1: Table S2). Whole genome sequence and whole genome genotype data were harmonized using the program PLINK [38], keeping only those sites common to all datasets and correcting SNP strand orientations as needed. A genotyping filter of 95% calls was applied to all populations.

### Global and local ancestry inference

Global continental ancestry estimates for each individual from the four LA populations were inferred using the program ADMIXTURE [15]. The harmonized SNP set was pruned using PLINK [38] with window size of 50bp, a step size of 10bp, and a linkage disequilibrium (LD) threshold of *r*^2^> 0.1, and ADMIXTURE was run with *K*=4 corresponding to African, European, Asian, and Native American ancestry components. Local continental ancestry estimates for each individual from the four LA populations were inferred using a modified version of the program RFMix [16] as previously described [39]. The chromosomal locations of ancestry-specific haplotypes were visualized with the program Tagore [40]. The complete harmonized SNP set was phased using the program SHAPEIT [41], and RFMix was run to assign African, European, or Native American ancestry to individual haplotypes from the LA populations.

### Single locus ancestry enrichment

Single gene (locus) ancestry enrichment (*z_anc_*) values were calculated for all three continental ancestry components (African, European, and Native American) across all four LA populations. Genomic locations of NCBI RefSeq gene models were taken from the UCSC Genome Brower (hg19 build) [42], and gene locations were mapped to the ancestry-specific haplotypes characterized using RFMix for each individual genome. For each gene, population-specific three-way ancestry fractions (*f_anc_*) were computed as the number of ancestry-specific haplotypes (ℎ_anc_), divided by the total number of ancestry-assigned haplotypes for that gene (*h*_*tot*_): *f_anc_* = *h_anc_*⁄*h*_*tot*_. Ancestry enrichment analysis was limited to genes that had *h*_*tot*_ values within one standard deviation of the genome-wide average for any population. Distributions of gene-specific ancestry fractions (*f_anc_*) for each population were used to calculate population-specific genome-wide average (*μ_anc_*) and standard deviation (*σ_anc_*) ancestry fractions. Then, for any given gene in any given population, ancestry enrichment (*z_anc_*) was calculated as the number of standard deviations above (or below) the genome-wide ancestry average: *z_anc_* = (*f_anc_* − *μ_anc_*)⁄*σ_anc_*, with gene-specific ancestry enrichment *P*-values computed using the *z* distribution. A Fisher’s combined score (*F*_*CS*_) was used to combine gene-specific ancestry enrichment *P*-values across the four LA populations as: 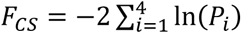. The statistical significance of *F*_*CS*_ was computed using the χ^2^ distribution with 8 (2*k*) degrees of freedom. Correction for multiple *F*_*CS*_ tests was performed using the Benjamini-Hochberg False Discovery Rate (FDR), with a significance threshold of *q* < 0.05 [43].

### Admixture simulation

Three-way admixed individuals were randomly simulated for each LA population – Colombia, Mexico, Peru, and Puerto Rico – and used to calculate expected levels of ancestry enrichment *z_anc_* as described in the previous section. Expected levels of *z_anc_* were combined across the four LA populations to yield expected Fisher’s combined scores (*F*_*CS*_) and their associated *P*-values as described in the previous section. Admixed populations were simulated as collections of genes (i.e. ancestry-specific haplotypes) randomly drawn from the genome-wide ancestry distributions for each LA population. Sized matched admixed populations were simulated for each LA population and combined to generate expected (*F*_*CS*_) and their associated *P*-values, and admixture simulation was also conducted across a range of population sizes (n=10 to 10,000) to evaluate the power of the combined evidence cross-population approach used to detect ancestry-enabled selection.

### Polygenic ancestry enrichment

Polygenic ancestry enrichment values (*PAE*) were computed by combining single locus ancestry enrichment values (*z_anc_*) across genes that function together to encode polygenic traits. Gene sets for polygenic traits were curated from a number of literature and database sources to represent a wide array of phenotypes (Additional file 1: Table S3). All gene sets were LD pruned with a threshold of r^2^ > 0.1 using PLINK. Additional details on the curation of polygenic trait gene sets can be found in Additional file 1 (page 12). For any trait trait-specific gene set, in any population, *PAE* was calculated by summing the gene-specific *z_anc_* values for all of the genes in the trait set: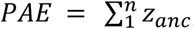, where *n* is the number of genes in the set. Since *z_anc_* values can be positive or negative, depending on over- or under-represented ancestry, values of *PAE* are expected to be randomly distributed around 0. The statistical significance levels of observed *PAE* values were calculated via comparison against distributions of expected *PAE* values calculated from 10,000 random permutations of size matched gene sets (Additional file 1: Figure S8). Observed values (*PAE_obs_*) were compared against the mean (*μ*_*PAE*_) and standard deviation (*σ*_*PAE*_) of the expected *PAE* values to compute the statistical significance for each trait: *z*_*PAE*_ = (*PAE_obs_*− *μ*_*PAE*_)⁄*σ*_*PAE*_, with *P*-values computed using the *z* distribution Correction for multiple tests was performed using the Benjamini-Hochberg False Discovery Rate (FDR), with a significance threshold of *q* < 0.05.

### Integrated Haplotype Scores (iHS)

Integrated Haplotype Scores (iHS) [21] were calculated for European and African continental populations from the 1KGP using the software selscan (version 1.1.0a) [44]. |iHS| scores were overlaid on genes with evidence of ancestry enrichment to scan for concurrent signals of selection.

### Modeling admixture-enabled selection

Admixture-enabled selection was modeled for the African enriched chromosome 6 MHC haplotype using a standard recursive population genetics model for positive selection [45]. Three allelic states were used for the selection model, each of which corresponds to a specific ancestry component: African, European, or Native American. Population-specific models were initialized with allele (ancestry) frequencies based on the genome-wide background ancestry fractions and run across a range of selection coefficient (*s*) values to determine the values of *s* that correspond to the observed African ancestry enrichment levels. This allowed us to compute a positive selection coefficient corresponding to the strength of African ancestry selection at the MHC locus for each population. Additional details of this model can be found in Additional file 1 (page 8-9 and Figure S7).

## Abbreviations

LA: Latin American
1KGP: 1000 Genomes Project
HGDP: Human Genome Diversity Project
SNP: single nucleotide polymorphism
iHS: integrated haplotype score

## Declarations

### Ethics approval and consent to participate

The de-identified human genome sequence data analyzed here are made publicly available and do not require ethics approval or consent to participate.

### Consent for publication

Not applicable

### Availability of data and materials

1000 Genomes Project (1KGP) data are available from http://www.internationalgenome.org/data/

Human Genome Diversity Project (HGDP) data are available from http://www.hagsc.org/hgdp/

Previously published Native American genotype data can be accessed from a data use agreement governed by the University of Antioquia as previously described [37].

### Competing interests

The authors declare that they have no competing interests.

### Funding

ETN, LR, ATC, ABC, and IKJ were supported by the IHRC-Georgia Tech Applied Bioinformatics Laboratory. KY was supported by the University of Georgia Research Foundation. AV-A was supported by Fulbright Colombia.

### Authors’ contributions

ETN conducted the ancestry enrichment analyses. LR performed the simulation analyses. ABC performed the genetic ancestry analyses. KY calculated the integrated haplotype scores. ETN, ATC, and AV-A curated the polygenic phenotype gene sets. ATC performed the polygenic phenotype significance testing. ETN and LR generated the manuscript figures. IKJ conceived of, designed, and supervised the project. ETN, LR, and IKJ wrote the manuscript. All authors read and approved the final manuscript.

## Additional files

Additional file 1: Figures S1-S8, Tables S2-S3, and Supplementary Methods

Additional file 2: Table S1

