## Additional File 1 for "Admixture-enabled selection for rapid adaptive evolution in the Americas"

Figure S1. Correspondence between continental ancestry estimates for LA populations generated by ADMIXTURE (x-axis) and RFMix (y-axis). 2

Figure S2. Comparison of global versus local continental ancestry inference for two admixed individuals. 3

Figure S3. Scheme of the ancestry enrichment analysis method used for this study. 4

Figure S4. Observed versus expected ancestry enrichment across the four LA populations studied here. 5

Figure S5. Ancestry enrichment power analysis. 6

Figure S6. African ancestry enrichment power analysis. 7

Modeling admixture-enabled selection at single loci 8

Figure S7. Modelling the strength of admixture-enabled selection at the MHC locus. 9

Table S2. Human populations analyzed as part of this study. 10

Polygenic ancestry enrichment 11

Table S3. Sources of the polygenic trait gene sets analyzed as part of this study. 11

Figure S8. Simulation of random polygenic trait gene sets used to compute statistical significance of polygenic ancestry enrichment ($PAE$). 12

References 13

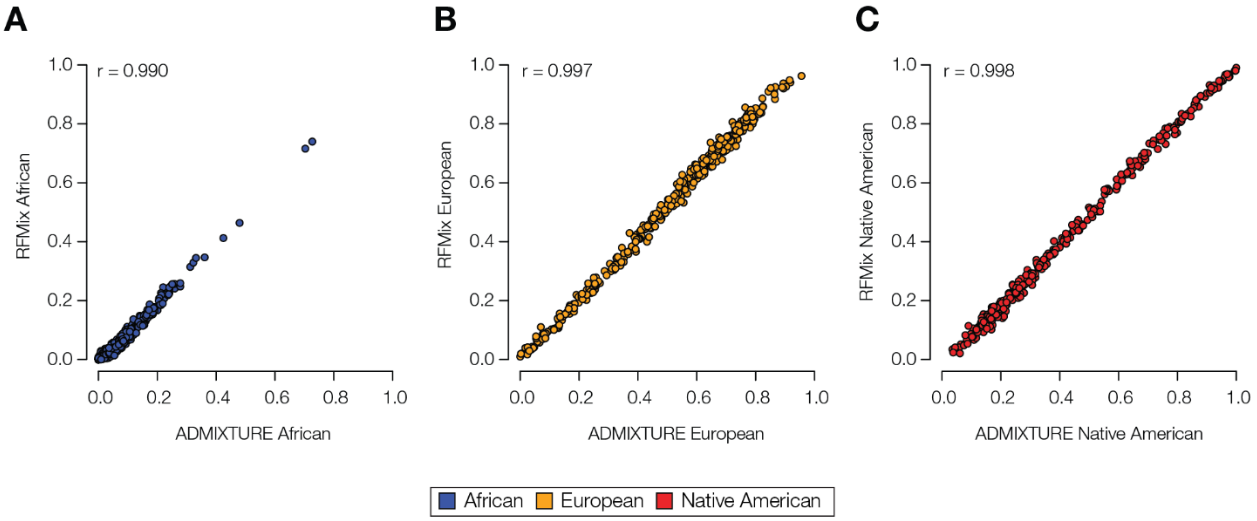


Figure S1. **Correspondence between continental ancestry estimates for LA populations generated by ADMIXTURE (x-axis) and RFMix (y-axis).** Ancestry estimates generated for individuals from the four LA populations studied here – Colombia, Mexico, Peru, and Puerto Rico – are shown separately for (A) African, (B) European, and (C) Native American ancestry. Correlation between ancestry estimates generated using the two programs are measured by Pearson's correlation coefficient (r).

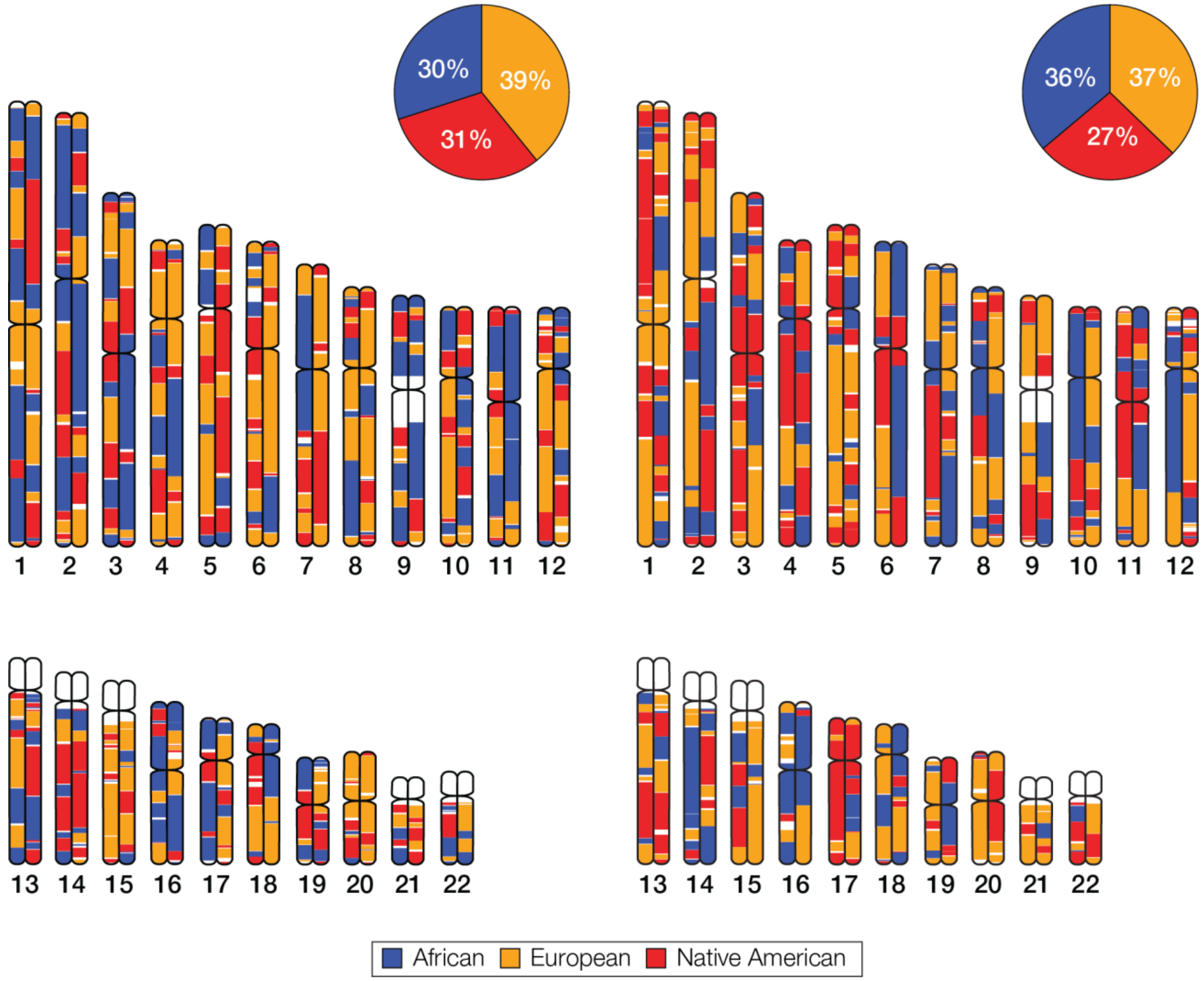


Figure S2. **Comparison of global versus local continental ancestry inference for two admixed individuals.** Continental ancestry estimates – for African (blue), European (orange), and Native American (red) ancestry components – were inferred using global (ADMIXTURE) and local (RFMix) ancestry methods. Global ancestry estimates are shown as pie charts, and local ancestry estimates (i.e. ancestry-specific haplotype assignments) are shown as chromosome ideograms. These two individuals have very similar levels of global ancestry but highly distinct local ancestry patterns.

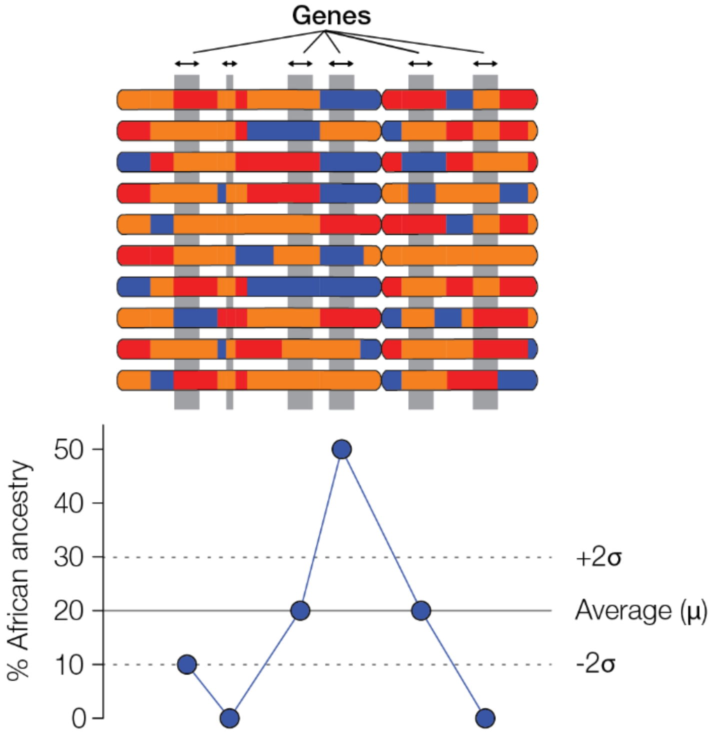


Figure S3. **Scheme of the ancestry enrichment analysis method used for this study.** Ten haploid chromosomes from a population are shown with corresponding regions aligned and ancestry-specific haplotypes indicated (as in Figure S2). Gene-specific ancestry enrichment ($z_{anc}$) is expressed as the number of standard deviations above or below the genome-wide ancestry fraction for the population.

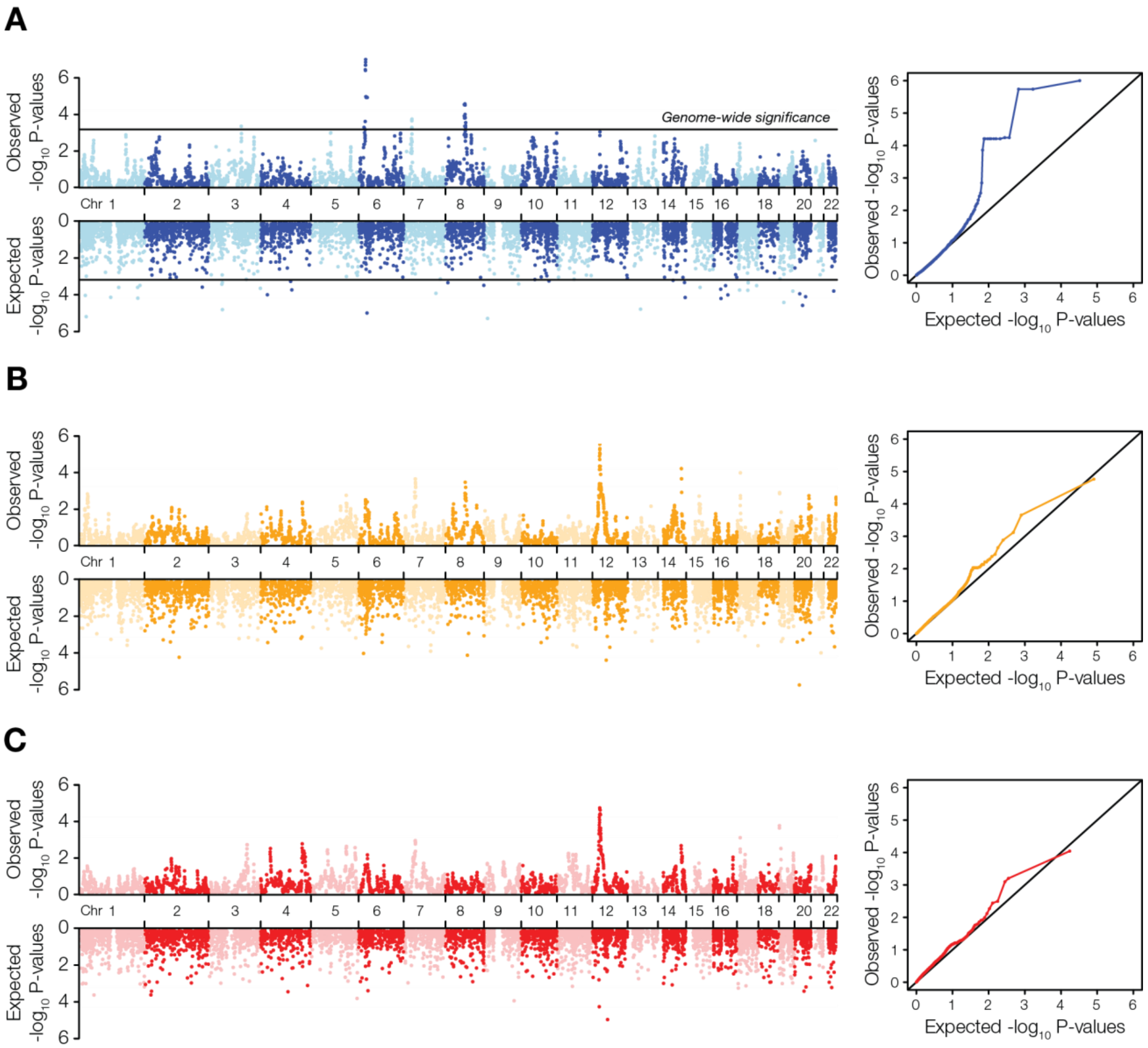


Figure S4. **Observed versus expected ancestry enrichment across the four LA populations studied here.** Manhattan plots and QQ plots show the genome-wide distributions of observed versus expected ancestry enrichment $-\log_{10} P$ values for African (A), European (B), and Native American (C) ancestry components. $P$-values correspond to the combined cross-population ancestry enrichment values ($F_{CS}$), calculated as described in the manuscript. Expected values were generated based on size matched simulated admixed populations, as described in the Methods section. On the African ancestry Manhattan plot, lines indicate genome-wide statistically significant cross-population ancestry enrichment (FDR $q < 0.05$). There were no genome-wide significant cross-population ancestry enrichment peaks for the European and Native American ancestry components. Ancestry-specific QQ plots also illustrate the deviation of observed versus expected ancestry enrichment for the African component, and the similarity of observed versus expected ancestry enrichment for the European and Native American components.

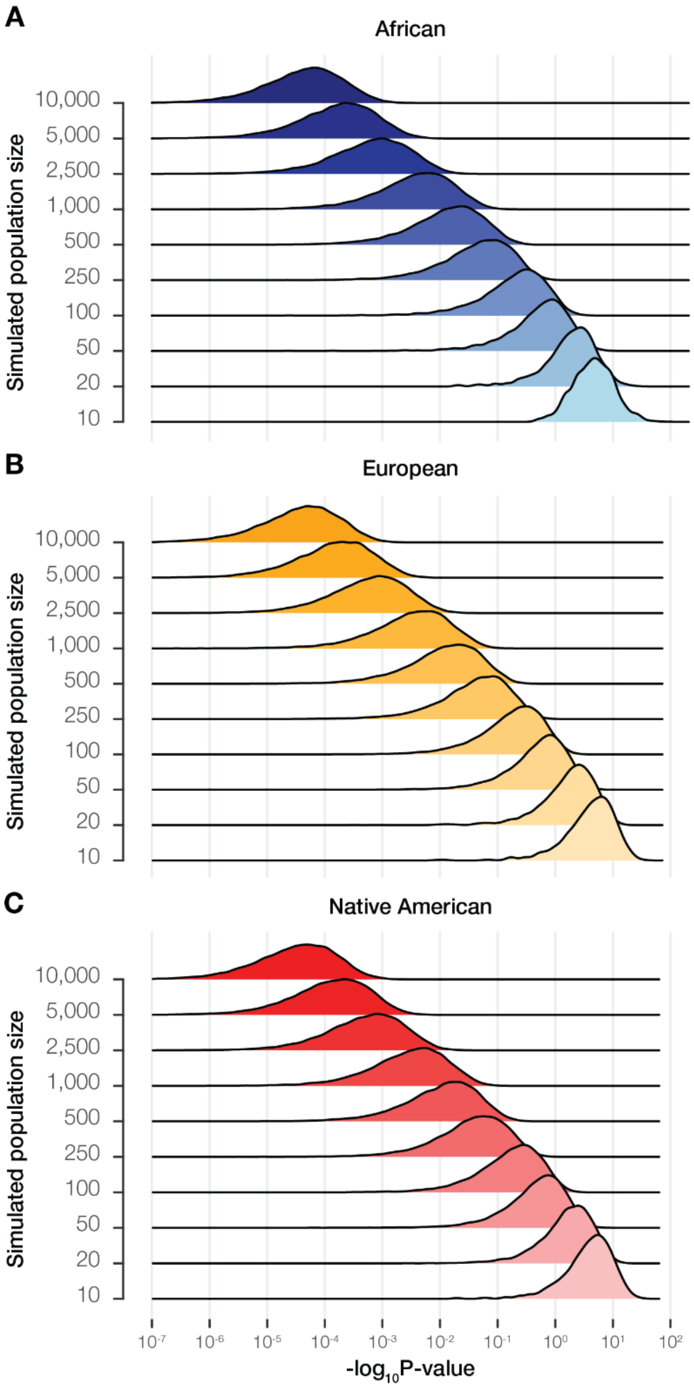


Figure S5. **Ancestry enrichment power analysis.** Four admixed LA populations were simulated across a range of population sizes (see Methods). Distributions of $-\log_{10} P$ values corresponding to combined cross-population ancestry enrichment values ($F_{CS}$) are shown across the range of simulated population sizes for each ancestry component: African (A), European (B), and Native American (C). This analysis was used as an additional, non-parametric method to compute the probability of observing ancestry-specific $F_{CS}$ by chance alone, at different population sizes.

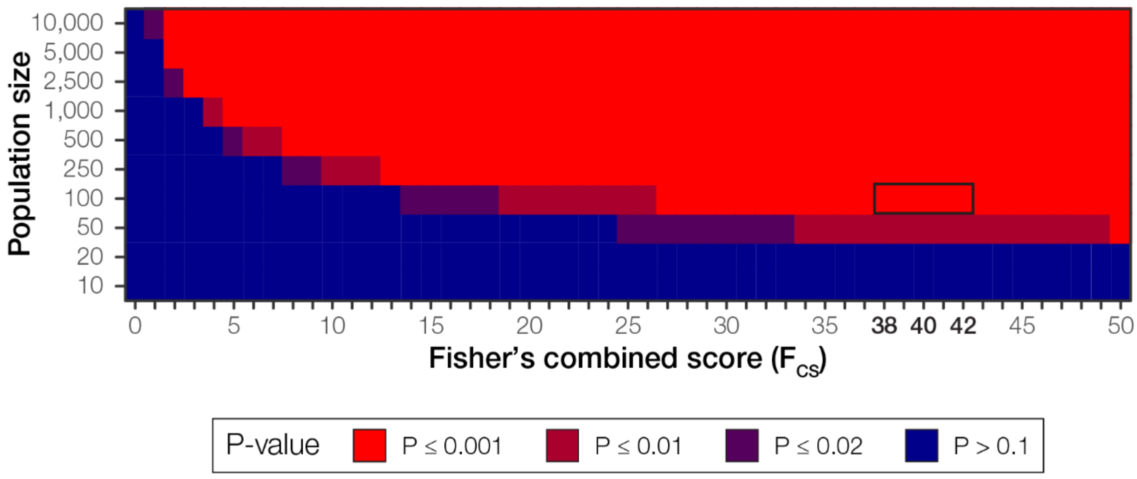


Figure S6. **African ancestry enrichment power analysis.** Four admixed LA populations were simulated across a range of population sizes (see Methods). Cross-population African enrichment values ($F_{CS}$), and their corresponding $P$ values, were computed across the range of population sizes (see Figure S5). The plot shows the relationship of $F_{CS}$ and $P$-values for different population sizes, i.e. the relationship between population size and the power of the cross-population ancestry enrichment metric. For African ancestry, a statistically significant FDR $q < 0.05$ corresponds to $P$<0.001 (bright red). Significant $P$ values are more common at higher sample sizes. The observed African $F_{CS}$ values at the MHC locus range from 38-42 (see boxed cells), which are well powered to detect significant ancestry enrichment at population sizes of ~100 individuals.

### Modeling admixture-enabled selection at single loci

A recursive triallelic model of positive selection was used to measure the strength of admixture-enabled selection at the chromosome 6 MHC locus in the Colombia, Mexico, and Peru populations. The model treats alleles as the three ancestry fractions: African, European, and Native American. This model is based on the approach to modelling selection used in the Populus software (https://cbs.umn.edu/populus/overview).

Assuming an African ancestry advantage, the relative fitness ($w_{ij}$) of a locus with ancestries $i$ and $j$ (ancestry genotypes) is calculated as:

$$w_{ij} = \begin{cases} 1, & if\ i = j = African \\ 1 - sh, & if\ i = African\ or\ j = African \\ 1 - s, & if\ i \neq African\ and\ j \neq African \end{cases}$$

where $s$ is the selection coefficient and $h$ is the dominance coefficient. This can be formulated as a fitness matrix:

| | | Ancestry $j$ | | |
|---|---|---|---|---|
| | | African ($A$) | European ($E$) | Native American ($N$) |
| Ancestry $i$ | African ($A$) | $1$ | $1-(sh)$ | $1-(sh)$ |
| | European ($E$) | $1-(sh)$ | $1-s$ | $1-s$ |
| | Native American ($N$) | $1-(sh)$ | $1-s$ | $1-s$ |

For each population, for each ancestry, the ancestry frequency in the next generation ($p_{i,t+1}$) was calculated as:

$$p_{i,t+1} = \frac{p_{i,t}w_i}{\bar{w}}$$

where $p_{i,t}$ is the ancestry frequency ($i$) in the current generation ($t$), $w_i$ is the marginal fitness of the ancestry, and $\bar{w}$ is the population mean fitness.

For each row of the fitness matrix, the marginal fitness ($w_i$) was calculated as

$$w_i = \sum_j w_{ij}p_j$$

where $w_{ij}$ is the relative fitness of the ancestry genotype and $w_{ij}$ is the frequency of the ancestry in the current generation.

For each generation, the population mean fitness ($\bar{w}$) was calculated as:

$$\bar{w} = \sum_i \sum_j w_{ij}p_ip_j$$

where $w_{ij}$ is the relative fitness of the ancestry genotype, $p_i$ is the frequency of the first ancestry, and $p_j$ is the second of the second ancestry.

For each population, the model was run using $s$ from 0-1.0, in increments of 0.001, and $h = 0.05$, until the final African ancestry frequency at 20 generations was the same as that observed for the chromosome 6 haplotype. The selection coefficient at this convergence was taken to be the strength of selection for the given population for the chromosome 6 African enriched haplotype.

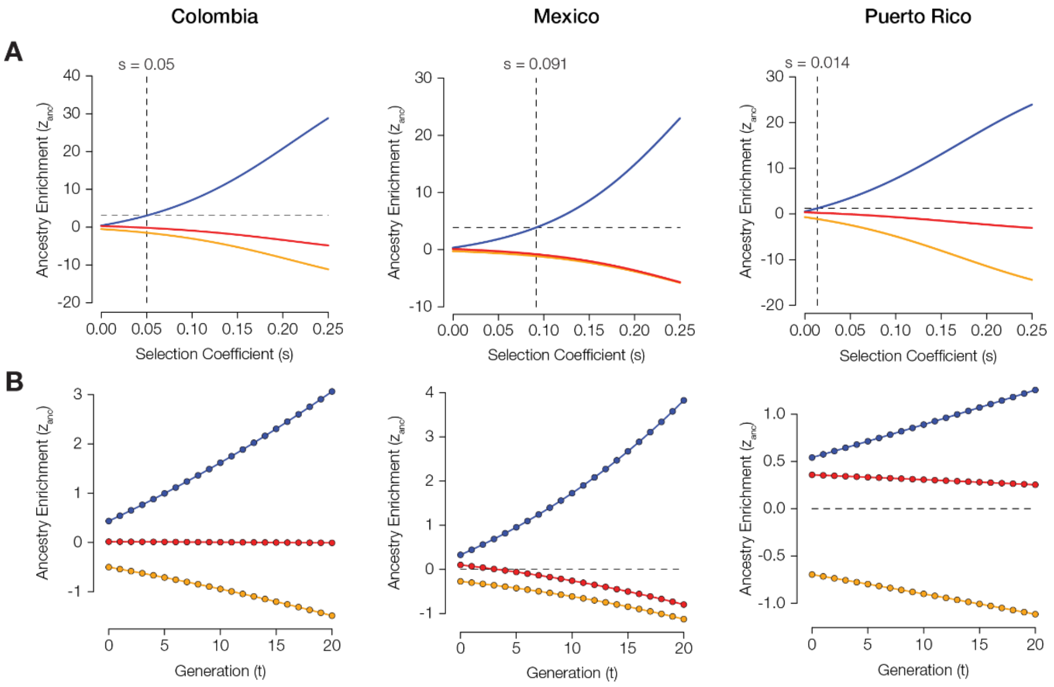


Figure S7. **Modelling the strength of admixture-enabled selection at the MHC locus.** The strength of admixture-enabled selection for African haplotypes at the MHC locus was modelled as described on pages 8-9. The three-ancestry population genetic model was run across a range of positive selection coefficient ($s$) values to identify the strength of selection needed to explain the observed levels of African ancestry enrichment at the MHC locus in the Colombia, Mexico, and Puerto Rico populations. The top panels for each population show the predicted levels of ancestry enrichment ($z_{anc}$, y-axis) across a range of different selection coefficients ($s$, x-axis). The intersection of the observed levels of African ancestry enrichment and their corresponding $s$-values are indicated with dashed lines. The bottom panels show the trajectory of predicted ancestry enrichment and depletion ($z_{anc}$, y-axis) over time ($t$ generations, x-axis) for each population given the inferred selection coefficient $s$ for each population.

**Table S2. Human populations analyzed as part of this study.** Populations are organized into continental groups, for both proxy ancestral reference populations and Latin American populations, and the number of genome (genotype) samples from each population is shown.

| Population group | Dataset[1] | Geographical source | n |
|---|---|---|---|
| African | 1KGP | Americans of African Ancestry in SW USA | 61 |
| | 1KGP | Mende in Sierra Leone | 85 |
| | 1KGP | African Caribbeans in Barbados | 96 |
| | 1KGP | Esan in Nigeria | 99 |
| | 1KGP | Luhya in Webuye, Kenya | 99 |
| | 1KGP | Yoruba in Ibadan, Nigeria | 108 |
| | 1KGP | Gambian in Western Division in the Gambia | 113 |
| European | 1KGP | British in England and Scotland | 91 |
| | 1KGP | Finnish in Finland | 99 |
| | 1KGP | Utah Residents (CEPH) with Northern and Western European Ancestry | 99 |
| | 1KGP | Iberian Population in Spain | 107 |
| | 1KGP | Toscani in Italia | 107 |
| Latin American | 1KGP | Mexican Ancestry in Los Angeles USA | 64 |
| | 1KGP | Peruvians from Lima, Peru | 85 |
| | 1KGP | Colombians from Medellin, Colombia | 94 |
| | 1KGP | Puerto Ricans from Puerto Rico | 104 |

| Population group | Dataset[1] | Geographical source | n |
|---|---|---|---|
| Native American | Reich et al | Waunana in Colombia | 3 |
| | Reich et al | Kogi in Colombia | 4 |
| | Reich et al | Mixtec in Mexico | 5 |
| | Reich et al | Embera in Colombia | 5 |
| | Reich et al | Guahibo in Colombia | 6 |
| | Reich et al | Ticuna in Brazil | 6 |
| | Reich et al | Guarani in Paraguay | 6 |
| | Reich et al | Piapoco in Colombia | 7 |
| | HGDP | Suruí in Brazil | 8 |
| | Reich et al | Inga in Colombia | 9 |
| | Reich et al | Wayuu in Colombia | 11 |
| | Reich et al | Kaqchikel in Guatemala | 13 |
| | HGDP | Pima in Mexico | 14 |
| | HGDP | Karitiana in Brazil | 14 |
| | Reich et al | Mixe in Mexico | 17 |
| | HGDP | Maya in Mexico | 21 |
| | Reich et al | Aymara in Bolivia | 23 |
| | Reich et al | Tepehuano in Mexico | 25 |
| | Reich et al | Quechua in Peru | 40 |
| | Reich et al | Zapotec in Mexico | 43 |

[1]1KGP = 1000 Genomes Project; HGDP = Human Genome Diversity Project; Reich et al [1]

**Polygenic ancestry enrichment**

**Polygenic trait gene set curation**

Gene sets for polygenic traits were curated from a number of literature and database sources, as shown below, to represent a wide array of phenotypes.

**Table S3. Sources of the polygenic trait gene sets analyzed as part of this study.** For each polygenic trait gene set source, the number of phenotypes and reference are shown.

| Phenotype source | n | Reference |
|---|---|---|
| NHGRI-EBI GWAS Catalog | 306 | [2] |
| InnateDB | 128 | [3] |
| Blood Transcription Modules | 209 | [4] |
| Blood Informative Transcripts | 9 | [5] |
| Innate Immune System | 9 | [6] |
| GIANT | 7 | [7] |
| Custom gene sets of interest | 10 | na |

Whenever possible, the gene sets were taken directly from the literature or the database. If gene sets were not directly accessible, SNP-level data was collected and directly mapped to genes to generate the gene set. Trait-specific gene sets from the NHGRI-EBI GWAS Catalog [2] were collected from SNP sets that were mapped using EBI's in-house pipeline; only SNPs that fell within a gene and were implicated at a genome-wide significance level of $P \leq 5x10^{-8}$ with the phenotype were used to generate the gene sets. Each polygenic trait from the Genetic Investigation of ANthropometric Traits (GIANT) consortium [7] had gene sets mapped according to specifications of the individual paper. All remaining gene sets from the literature were mapped using NCBI's dbSNP database as needed. To control for any linkage between genes, linkage disequilibrium (LD) pruning was performed on each gene set using PLINK. For each set, pruning was performed on all gene pairs with genic SNP $r^2 > 0.1$; if this was found to be true then only one member of the gene pair was retained for polygenic ancestry enrichment analysis. After LD pruning was complete, the gene sets were filtered based on size so that all phenotypes included in the analysis contained two or more genes.

**Statistical significance of polygenic ancestry enrichment**

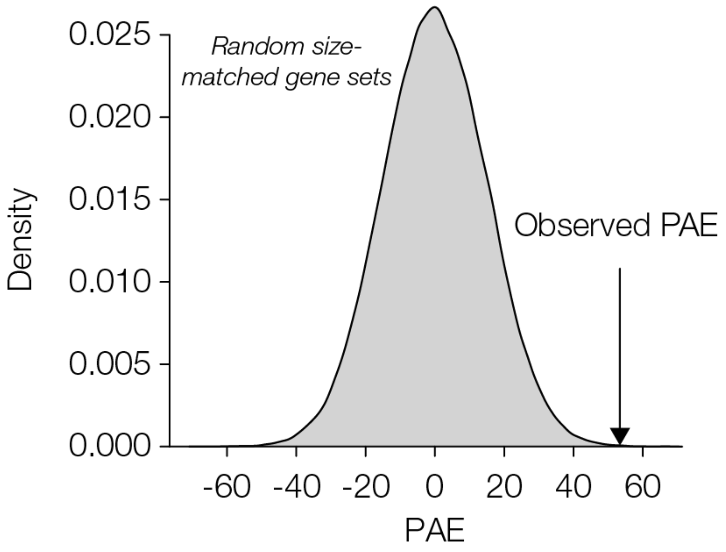


Figure S8. **Simulation of random polygenic trait gene sets used to compute statistical significance of polygenic ancestry enrichment ($PAE$).** Size-matched random gene sets are simulated to generate an expected (null) distribution of $PAE$ values.
